# Long-term influence of climate and experimental eutrophication regimes on phytoplankton blooms

**DOI:** 10.1101/658799

**Authors:** Kateri R. Salk, Jason J. Venkiteswaran, Raoul-Marie Couture, Scott N. Higgins, Michael J. Paterson, Sherry L. Schiff

## Abstract

Phytoplankton blooms respond to multiple drivers, including climate change and nutrient loading. Here we examine a long-term dataset from Lake 227, a site exposed to a fertilization experiment (1969–present). Changes in nitrogen:phosphorus loading ratios (high N:P, low N:P, P-only) did not impact mean annual biomass, but blooms exhibited substantial inter- and intra-annual variability. We used a process-oriented lake model, MyLake, to successfully reproduce lake physics over 48 years and test if a P-limited model structure predicted blooms. The timing and magnitude of blooms was reproduced during the P-only period but not for the high and low N:P periods, perhaps due to N acquisition pathways not currently included in the model. A model scenario with no experimental fertilization confirmed P loading is the major driver of blooms, while a scenario that removed climate-driven temperature trends showed that increased spring temperatures have exacerbated blooms beyond the effects of fertilization alone.

**Significance Statement:** Harmful algal blooms and eutrophication are key water quality issues worldwide. Managing algal blooms is often difficult because multiple drivers, such as climate change and nutrient loading, act concurrently and potentially synergistically. Long-term datasets and simulation models allow us to parse the effects of interacting drivers of blooms. The performance of our model depended on the ratio of nitrogen to phosphorus inputs, suggesting that complex biological dynamics control blooms under variable nutrient loads. We found that blooms were dampened under a “no climate change” scenario, suggesting that the interaction of nutrient loading and increased temperature intensifies blooms. Our results highlight successes and gaps in our ability to model blooms, helping to establish future management recommendations.

**Data Availability Statement:** Data and metadata will be made available in a GitHub repository (https://github.com/biogeochemistry/Lake-227). Upon manuscript acceptance, the repository will be made publicly available and a DOI will be provided. We request that data users contact the Experimental Lakes Area directly, per their data use policy (http://www.iisd.org/ela/wp-content/uploads/2016/04/Data-Terms-And-Conditions.pdf).

## Introduction

Eutrophication and harmful algal blooms (HABs) are global water quality issues, increasing in occurrence and intensity in lakes worldwide (Heisler et al. 2008; Paerl et al. 2018). Anthropogenic nutrient loading is the major driver of excess phytoplankton growth (Carpenter et al. 1998), and climate-related drivers including temperature and precipitation are additional influencing factors (Paerl et al. 2011; Sinha et al. 2017). Changes in these external drivers can induce regime shifts in lakes, which often occur abruptly (Ratajczak et al. 2018). Mitigating the negative effects of eutrophication and HABs requires a nuanced understanding of the interacting effects of external drivers (Elliott 2012).

Whole-ecosystem experiments have demonstrated their potential to reveal the effect of nutrient loading on lake eutrophication (Schindler 1998). Lake 227 at the IISD-Experimental Lakes Area (IISD-ELA; Ontario, Canada) is a well-known example of one such whole-ecosystem manipulation. The experiment has been conducted for five decades, during which artificial N and P loadings were added in molar ratios of 27:1 (1969-1974), 9:1 (1975-1989), and 0:1 (P-only, 1990-present) while P loads were held constant. These abrupt transitions in external nutrient loads and stoichiometric ratios enable examination of phytoplankton responses to changing fertilization. Decreases in N loading in Lake 227 resulted in the dominance of diazotrophic cyanobacteria during the mid-summer phytoplankton bloom (Hendzel et al. 1994), whereas annual biomass remained unchanged due to a constant P supply (Schindler et al. 2008; Paterson et al. 2011; Higgins et al. 2017). Lake 227 offers a unique opportunity to leverage a five-decade fertilization experiment to test our ability to characterize the drivers of phytoplankton blooms and examine the role of climate on interannual variation in biomass.

The response of lakes to concurrent drivers of eutrophication can be explored using process-oriented models (Couture et al. 2018; Page et al. 2018; Janssen et al. 2019). A variety of lake ecosystem models exist that include physical processes and nutrient dynamics, varying in modeling approach, spatial dimensions, and complexity of process representation (Robson 2014). Models using a one-dimensional (1D) physical driver have been successfully applied to small lakes where horizontal mixing is rapid (Saloranta and Andersen 2007). Given the tradeoffs in choosing an appropriate model for a given system, 1D models strike a balance between computation effort, data requirements, and ability to determine the extent to which physical drivers will influence the nutrient cycling and phytoplankton growth over multi-year timescales.

Here, we explore inter- and intra-annual bloom dynamics, using statistical approaches and process-oriented modeling to integrate climatic variables and fertilization regime as drivers of phytoplankton growth. Specifically, we aim to (1) determine if the hypothesis of P-limitation appropriately reproduces bloom dynamics across different fertilization regimes, and (2) disentangle the concurrent effects of climate and nutrient loading on phytoplankton blooms over five decades.

## Methods

### I. Historical monitoring data

Lake 227 is a small (5 ha) headwater lake in northwest Ontario, Canada. The hydrology of Lake 227 is relatively simple, with a bowl-shaped morphometry (mean depth 4.4 m, maximum depth 10 m) and negligible groundwater inputs. Starting in 1969 and continuing through present, physical, chemical, and biological data were collected biweekly in Lake 227 during the ice-free season and 2-4 times during the ice-covered period. Meteorological data, including air temperature, wind speed, precipitation, and radiation, were collected at a station jointly operated with Environment and Climate Change Canada, located 4.1 km from Lake 227. Loads of nutrients from the Lake 227 watershed were estimated using data from streams flowing into a nearby reference lake, Lake 239, which were monitored continuously for flow and weekly for nutrient chemistry during the ice-free season. Measurements of wind speed, radiation, ice phenology, and inflow volume and temperature were scaled to the conditions at Lake 227 (Supplemental Information).

Statistical analyses of climate, nutrient, and phytoplankton data were conducted to determine trends in drivers and effects over time. Meteorological datasets included year-round measurements, whereas lake and stream datasets included measurements only from the ice-free season. Trends in lake and stream data over time were determined by a Mann-Kendall test with the Yue and Wang (2004) variance correction approach to account for the presence of autocorrelation in the data (Supplemental Information). Potential change points in each dataset were determined by Pettitt’s test. Trends in daily meteorological data over time were determined by a Seasonal Mann-Kendall test. All statistical analyses and data visualization were carried out in R version 3.5.1 (R Core Team 2013). Code and data available from [placeholder data citation].

### II. Lake modeling

MyLake is a 1D process-oriented model designed to simulate seasonal ice and snow cover, heat exchange and thermal stratification, nutrient cycling, phytoplankton growth, and water-sediment coupling (Saloranta and Andersen 2007; Supplementary Information). MyLake has been updated with oxygen and DOC dynamics (Couture et al. 2015; de Wit et al. 2018). The model has previously been applied to boreal lakes to simulate the response of P cycling to external P loading and climate change (Couture et al. 2014, 2018). MyLake does not include top-down controls on phytoplankton biomass.

The MyLake model was chosen for Lake 227 owing to its ability to characterize the thermal structure, catchment inputs, and biogeochemical reactions in the water column and sediment. The rationale for choosing a P-limited phytoplankton growth model was supported from previous studies demonstrating that P was the main driver of phytoplankton biomass in Lake 227 (Paterson et al. 2011; Higgins et al. 2017). Prior work suggests that the filamentous phytoplankton community was generally not a high quality food source for zooplankton in Lake 227, which were dominated by rotifers (Paterson et al. 2011). While grazing can occasionally be important in Lake 227 (Elser et al. 2000; Paterson et al. 2002), we prioritized the reproduction of physical and biogeochemical aspects of the system over food web interactions.

We simulated ice formation, water temperature, dissolved oxygen and phytoplankton bloom dynamics in Lake 227 from 1969 to 2016. Particulate P (PP) was chosen as a proxy for phytoplankton biomass to assess model performance because the vast majority of PP in Lake 227 is present in phytoplankton tissues rather than bacterial or allochthonous pools (Hecky et al. 1993; Elser et al. 1995). While biomass and chlorophyll are also measured in Lake 227, these metrics are not state-variables of the MyLake model and would require conversion factors that can vary considerably *in situ* (Hecky et al. 1993). Alternatively, PP is measured in Lake 227 and modeled directly by MyLake, enabling comparisons between observed and modeled concentrations.

Application of MyLake to Lake 227 involved a characterization of initial water column and sediment conditions, inputs of daily meteorological conditions and inflow chemistry, and parameterizations for physical, chemical, and biological reactions (Supplementary Information). Parameters for PP- and TDP-sensitive parameters were optimized using the Genetic Algorithm function of the MATLAB Global Optimization Toolbox, which seeks to minimize the sum of squared error between observed and modeled PP and TDP by varying parameters within the maximum probable range. Based on changes in nutrient stoichiometry and shifts in phytoplankton communities with different growth and nutrient uptake characteristics, parameterizing the model separately for the three periods was deemed appropriate. Optimization was performed for the first five years in each fertilization period, and performance was validated across the entire period. 1969 was designated as a model spin-up year and was excluded from final analyses. Model output from 1996 was also left out of post-processing analyses due to a trophic cascade experiment that resulted in unusually high zooplankton grazing in that year (Elser et al. 2000). Model code, input and output files are available from [placeholder data citation].

The goodness of fit between modeled and observed values in Lake 227 was assessed with several metrics. The root-mean squared error (RMSE) quantified the average deviation of modeled values from observed values in the units of interest. Cumulative PP was calculated as the sum of epilimnion PP concentrations between 16 May and 31 October each year, sourced from daily output (modeled) and estimated by linear interpolation between monitored dates (observed). Model performance for O_2_ was evaluated at 4 m depth, which was typically associated with the metalimnion during the stratified period and experiences dynamic physical and biological conditions. Aggregated performance metrics for temperature, PP, TDP, and O_2_ were calculated in terms of normalized bias (*B*\*) and normalized unbiased root mean squared difference (*RMSD*’*; Supplementary Info; Los and Blaas 2010). *B*\* described a systematic over- or underestimation of the modeled values. *RMSD*’* represented the difference in standard deviation between modeled and observed values, with negative values indicating lower variance in the observed values than the modeled values and vice versa. Values of *B*\* and *RMSD*’* were plotted in a target diagram (Taylor 2001), allowing for comparisons of model performance among the three fertilization periods independently of the magnitude of each variable.

Finally, to separate the potential effects of climate change and fertilization on phytoplankton blooms in the P-only period (1990–2016), three model scenarios were run. The first scenario included observed air temperature and external fertilization (Climate Change + Fertilization), representing actual historical conditions. The second scenario included observed air temperature but excluded external fertilization (Climate Change Only). The third scenario included external fertilization and a detrended air temperature input (Fertilization Only), computed using a loess-based time series decomposition on a 365-day seasonal cycle. This detrending approach preserved seasonal and random variability in temperature but removed the systematic increase in temperature since 1969 (3.2 ± 1.3 °C, range 0.7–6.5; Figure S1, S2).

## Results

### I. Historical Monitoring Data

In response to a decrease in fertilizer N:P ratios in 1975 and 1990 (Figure 1a), diazotrophic cyanobacteria (*Aphanizomenon* spp.) became increasingly dominant during the summer phytoplankton bloom (Figure 1b). Epilimnetic TN:TP molar concentrations increased until 1996, then remained steady through the P-only fertilization period (Figure 1c, Table S1). Epilimnetic TDN:TDP molar concentrations increased until 1975, then decreased significantly through the low N:P and P-only fertilization periods (Figure 1d, Table S1). PP concentrations were steady from the beginning of the high N:P fertilization period until 1978, then decreased significantly through the low N:P and P-only fertilization periods (Figure 1e, Table S1). Chl concentrations displayed a significant decreasing trend both before and after the change point, which occurred in 1987 (Figure 1f, Table S1).

**Figure 1.**
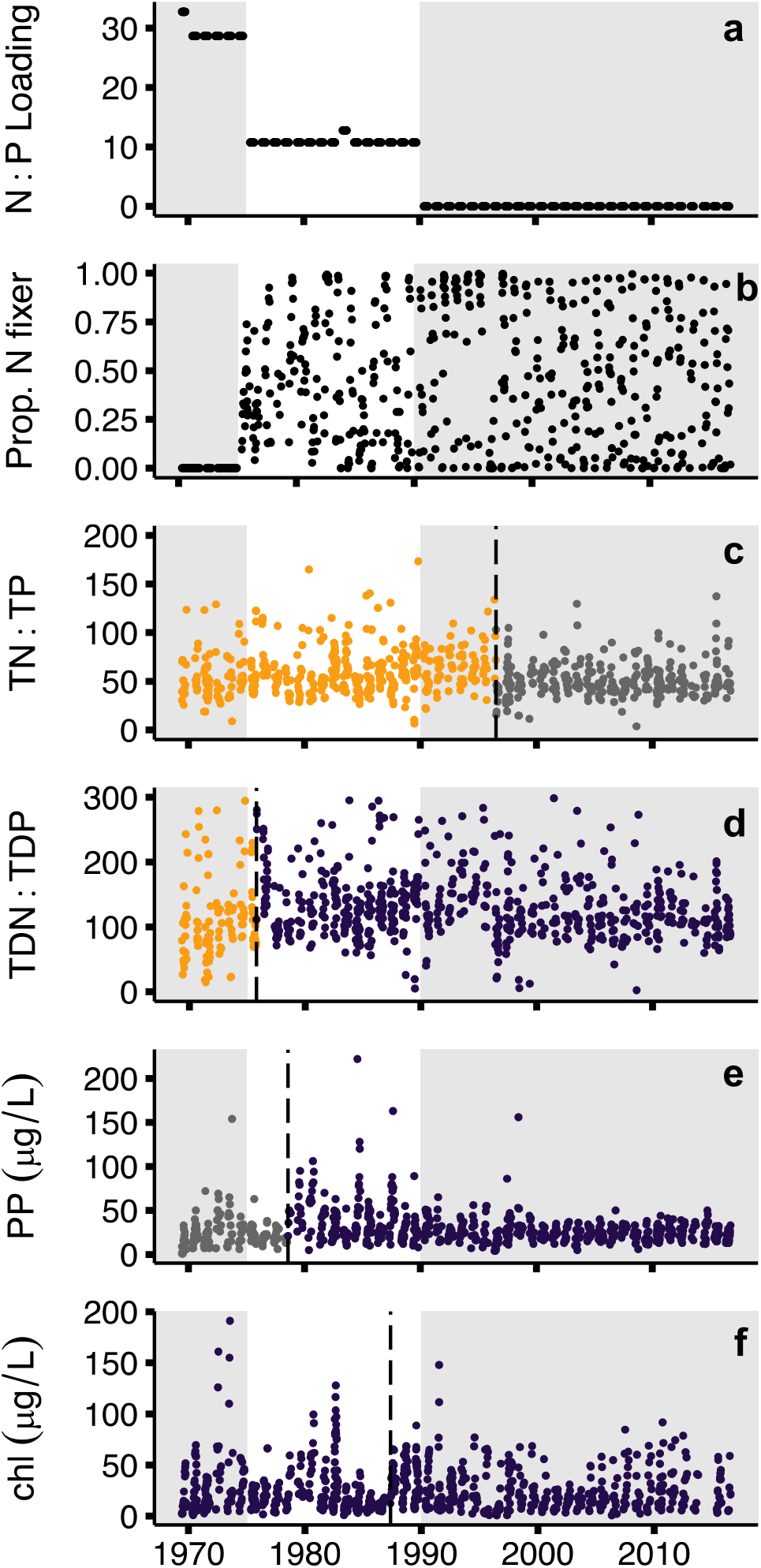
Observed historical trends in the epilimnion of Lake 227 for the ice-free season. Points represent twice-monthly data collections. Shadings represent different N:P fertilization periods. Dotted line represents breakpoint as determined by Pettitt’s test. Color of point indicates direction of trend before or after breakpoint as indicated by Mann-Kendall test (yellow = significant positive trend, purple = significant negative trend, gray = non-significant trend). (a) Molar ratios of fertilizer N:P. (b) Proportion of phytoplankton cells identified as diazotrophic. (c) and (d) *In situ* molar ratios of total and dissolved N:P, respectively. (e) and (f) *In situ* concentrations of particulate phosphorus and chlorophyll *a*.

Climatic drivers displayed substantial inter- and intra-annual variation (Figure S3, Table S1, S2). A Seasonal Mann-Kendall test indicated a significant increase in overall air temperature over the 48-year period, while the total temperature range has narrowed. Wind speed and daily radiation decreased significantly over time. Precipitation increased significantly, with largest intensity rain events occurring after 1980. Catchment inflows of P and N were generally a small fraction of fertilizer inputs, with TP inflow concentrations decreasing significantly over time and DIN concentrations displaying no monotonic trend.

### II. Assessment of Model Fit

Across the 48-year period, ice break and ice freeze dates were predicted within 6 days or less. Modeled ice phenology closely followed the variation experienced in Lake 239, indicating the model responded to local climatic drivers appropriately (Figure 2a). Modeled water temperatures followed the seasonal trends of warming and cooling observed in Lake 227 (Figure 2b). Temperatures at 1, 4, and 9 m depth were predicted within 1.92, 1.65, and 0.48 °C, respectively. Model performance for the thermal structure of the lake was similar across fertilization periods (Table S3).

**Figure 2.**
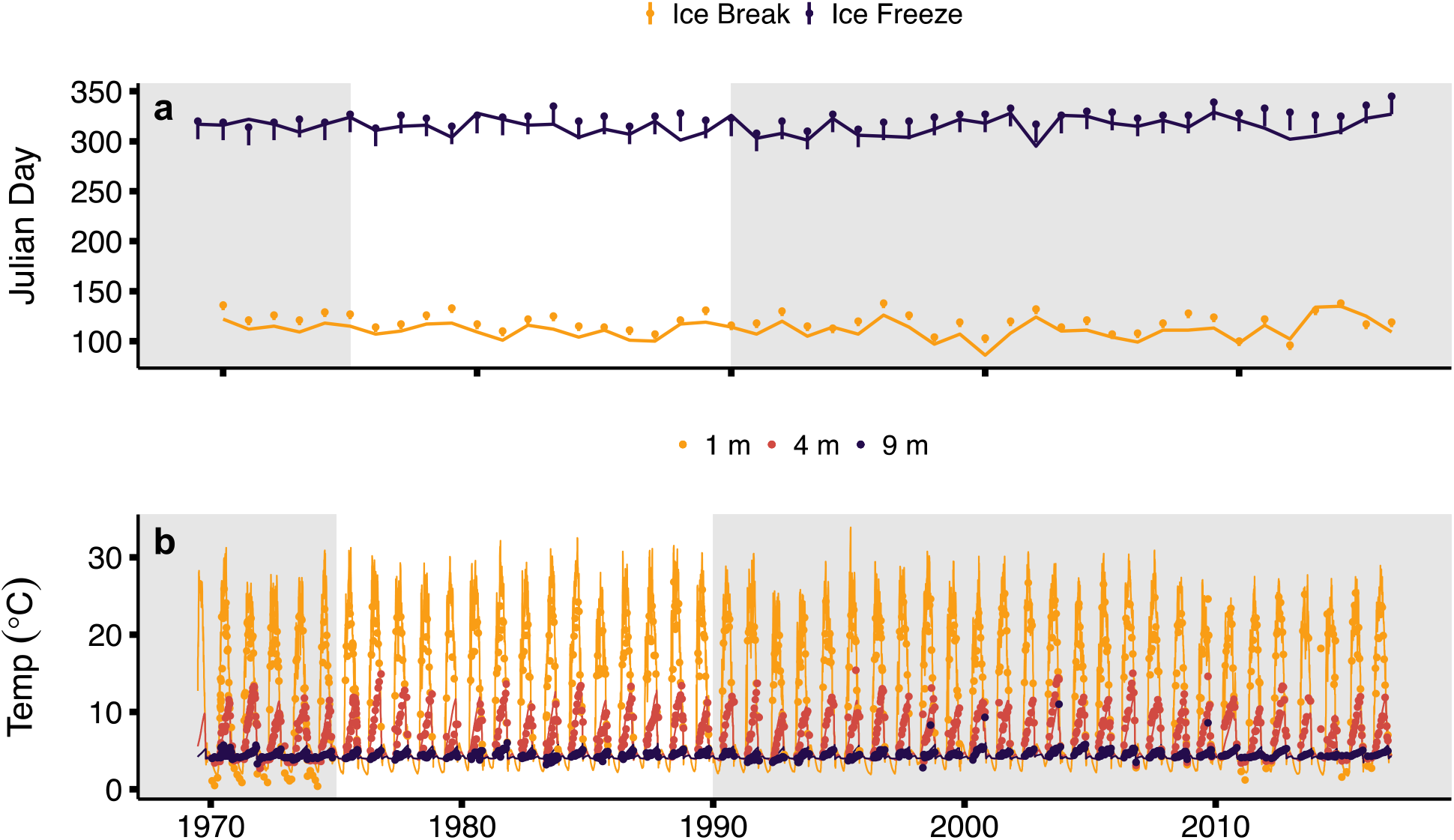
Physical model fit for three N:P fertilization periods. (a) Ice break and ice freeze dates, as predicted by the model (lines) and as observed in Lake 239 (dots). Tails on dots represent the range in ice break and freeze dates predicted for Lake 227 due to its size, up to five and 18 days earlier than Lake 239, respectively. (b) Temperature at 1, 4, and 9 m depth as predicted by the model (lines) and as observed in Lake 227 (dots). Shadings represent different N:P fertilization periods.

The fertilization periods displayed differences in phytoplankton communities and the range of epilimnetic PP concentrations (Figure 1a, e). To reproduce these patterns, optimizations of PP- and TDP-sensitive parameters yielded different parameter sets for the three periods, namely for the half saturation growth P level (P_half; Table S4). The model predicted a fairly consistent annual peak and cumulative PP concentration, in contrast to observations that displayed substantial interannual variation (Figure 3). RMSE for PP was 20.6 and 21.8 μg/L for the high and low N:P periods, respectively. For the P-only period, RMSE for PP was 10.5 μg/L, representing a doubling in the accuracy of model predictions compared to the earlier fertilization periods. During the P-only period, the timing and magnitude of the phytoplankton bloom were captured by the model for both PP and TDP concentrations (Figure S4).

**Figure 3.**
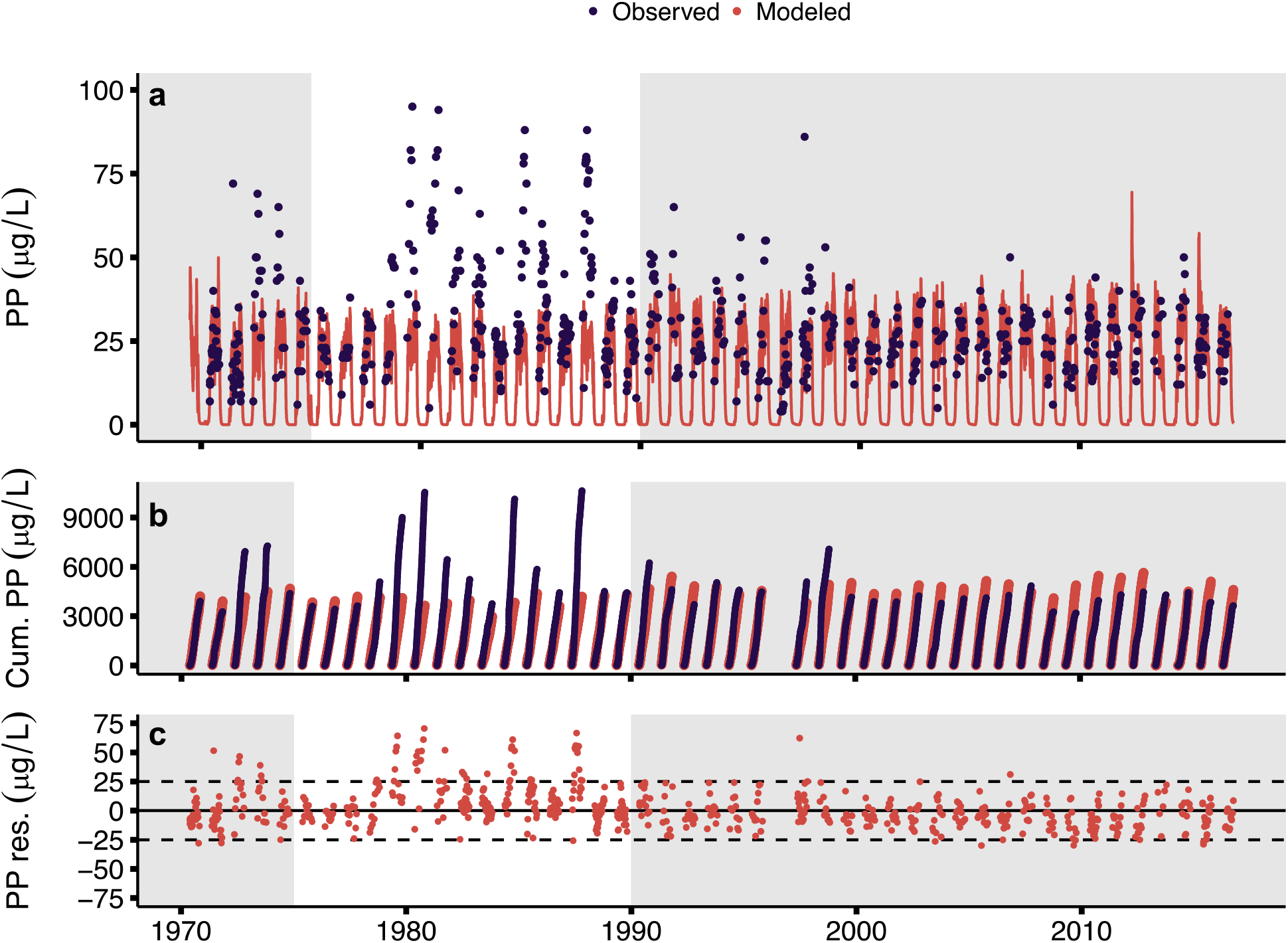
Model performance for particulate phosphorus (PP). (a) PP concentrations observed in Lake 227 (dots) and as predicted by the model (red lines). (b) Observed (purple) and modeled (red) cumulative PP, as calculated by the sum of daily concentrations in the ice-free season. (c) PP residuals (modeled PP – observed PP). Shadings represent different N:P fertilization periods.

Model performance for temperature at 1, 4, and 9 m depth was within one unit of *B*\* and *RMSD*’* for all fertilization periods (Figure 4). O_2_ at 4 m depth was modeled with similar success for the three periods. For P partitioning, the model performed best for the P-only period, with PP and TDP values for *B*\* of 0.37 and −0.36, respectively, and *RMSD*’* of −3.57 and 6.5, respectively. For the high and low N:P periods, PP and TDP were within one unit of *B*\*, but variance was greatly underestimated for PP (*RMSD*’* of −15.4 and −15.9, respectively) and greatly overestimated for TDP (*RMSD*’* of 22.1 and 8.7, respectively).

**Figure 4.**
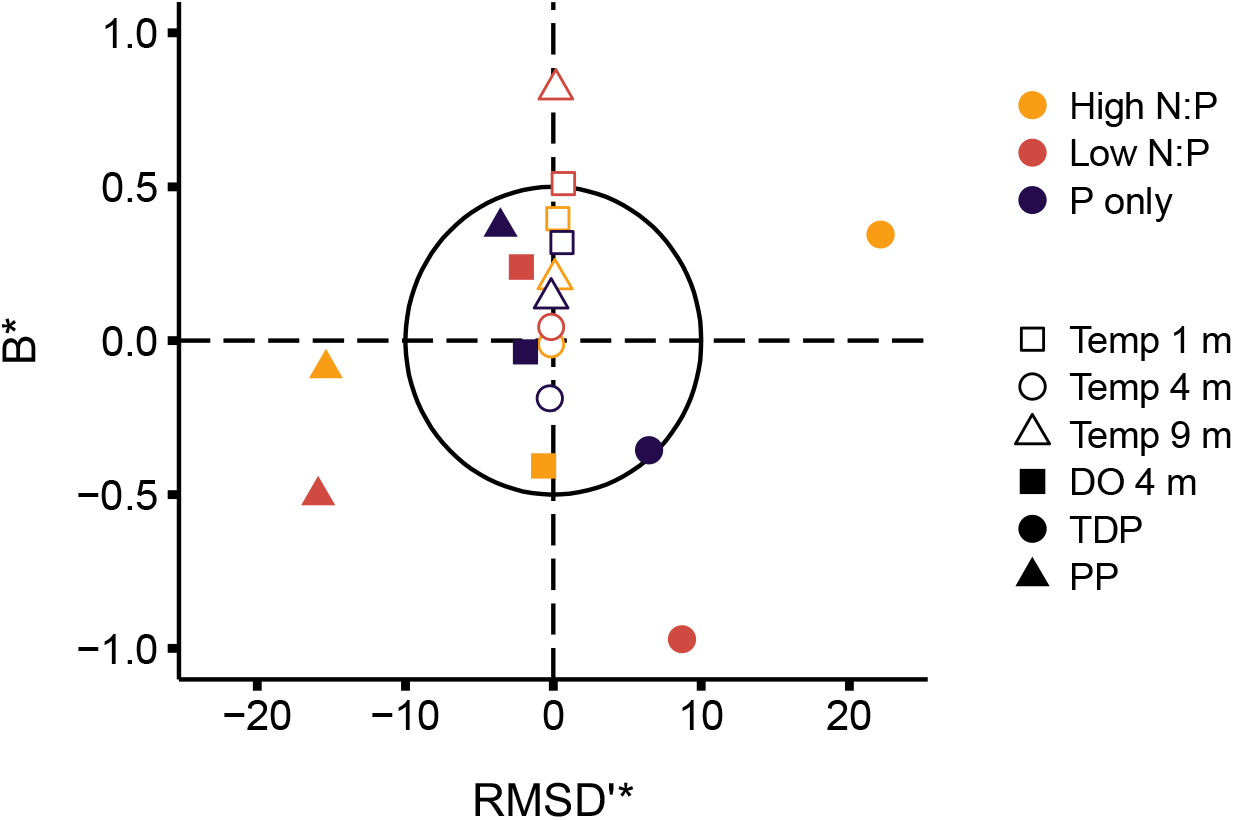
Target plot evaluating model fit for three periods. Values of normalized unbiased root mean squared difference (*RMSD*’*) and normalized bias (*B*\*) near zero represent a model fit with similar variance and minimal over- or underestimation of values compared to observations, respectively. The circle represents absolute values of *RMSD*’* and *B*\* of 10 and 0.5, respectively.

Scenario analysis was deemed appropriate for the P-only period, as the physical aspects of the model tracked historical observations and the timing and magnitude of PP concentrations were reproduced. Detrended air temperature in the Fertilization Only scenario, relative to 1969, resulted in shifts in surface water temperature, with the greatest difference occurring in April and May (Figure S5). Daily comparisons of modeled PP concentrations showed a significant difference among scenarios (paired Wilcoxon test, p < 0.0001; Figure 5a). Daily PP concentrations for the ice-free season in the Climate Change + Fertilization scenario exceeded those in the Fertilization Only scenario 87 % of the time, with the greatest differences occurring from June to September and the smallest differences in May and October (Figure 5b). Peak annual PP concentrations differed significantly among scenarios (Kruskall-Wallis test; X^2^ = 56.14, p < 0.0001) and were greatest in the Climate Change + Fertilization scenario (42.3 ± 7.1 μg/L, range 31.9–69.5), intermediate in the Fertilization Only scenario (37.5 ± 5.8 μg/L, range 27.4–47.9), and lowest in the Climate Change Only scenario (1.6 ± 0.9 μg/L, range 0.4–3.9). Peaks in PP concentration occurred in the early part of the ice-free season in the Climate Change Only scenario and generally occurred in the late part of the ice-free season in the Fertilization Only scenario (Figure 5a).

**Figure 5.**
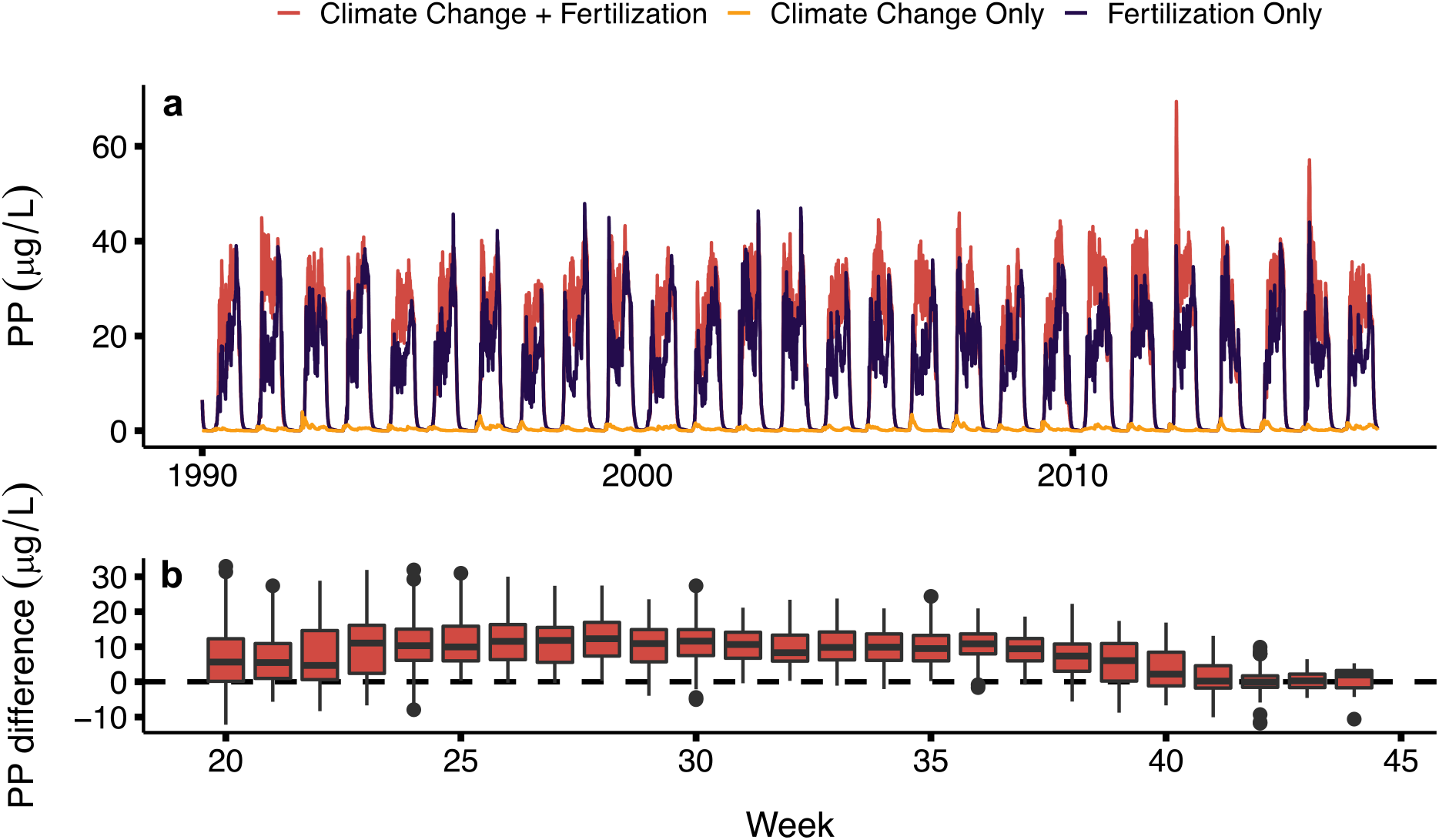
Climate and fertilization scenario analysis for the P-only regime (1990-2016). (a) Daily PP concentrations for the Climate Change + Fertilization (i.e., actual historical conditions; red), Climate Change Only (i.e., no external P loading; yellow), and Fertilization Only (i.e., historical P loading and detrended temperature; purple) model scenarios. (b) The difference in daily PP measurements between the Climate Change + Fertilization and Fertilization Only scenarios, pooled for each week across modeled years (1990-2016). Positive values indicate higher PP in the Climate Change + Fertilization scenario than in the Fertilization Only scenario.

## Discussion

Lake 227 has responded to shifts in N and P loading through changes in nutrient stoichiometry, phytoplankton community composition, and biomass. Although ratios of TDN:TDP in Lake 227 decreased concomitant with a reduction in external N fertilization, biological N fixation by cyanobacterial diazotrophs maintained TN:TP ratios across the fertilization regimes. The shift from an increasing trend in TN:TP to a steady ratio of TN:TP occurred in 1996, potentially due to a lag time coinciding with the 7-year hydraulic residence time of the lake or a trophic cascade experiment that left the lake fishless and potentially altered recycling pathways of N and P (Elser et al. 2000). While previous analyses have indicated that Lake 227 has not experienced a decline in annual mean epilimnetic phytoplankton biomass indices (Paterson et al. 2011), the time series analysis incorporating all measurements in the ice-free season revealed shifts in interannual bloom composition occurring over decades. The decreasing trend of chl and PP concentrations moving from the low N:P to P-only fertilization regime highlights that blooms have a more consistent lower peak in the P-only period compared to previous periods, and PP and chl concentrations above 100 μg/L are no longer observed. Since no concomitant downward trend in annual mean biomass has occurred, this result suggests that blooms have become lower in peak magnitude but spread out over a greater period in the ice-free season in the P-only period, in contrast to the high and low N:P periods with more intense yet short-lived blooms.

Our model application illuminates the complexity associated with phytoplankton blooms and the ongoing challenge of applying process-oriented ecosystem models to characterize seemingly well-understood systems. Lake 227 is small, bowl-shaped lake with a well-known fertilization regime and long-term monitoring record, providing a “best case” to examine model performance. Many systems experience dramatic shifts in stoichiometry and nutrient loading (Jeppesen et al. 2005; Hessen et al. 2009), inducing changes in lake ecosystem functioning that are difficult to capture by the fixed structures of process-oriented models. This challenge is exemplified here by variable model performance with respect to fertilization regime. The best performance indicators of bias and variance for PP and TDP were observed in the P-only period, whereas the variance in PP was substantially underestimated and the concentrations and variance in TDP were overestimated during the high and low N:P periods (Figure 4). The high and low N:P periods experienced both the highest and the lowest peak and cumulative PP concentrations, but this variability was not captured by the model (Figure 3). Given that nutrient additions were well-constrained and annual P additions were consistent over the study period, the large swings in observed PP concentrations among years are unexpected. Evidently, there are sources of variability introduced during the high and low N:P regimes that are unaccounted for by the model structure. Because of the excellent performance of the hydrodynamics component of the model, this source of variability is more likely in the biological component, namely a suite of processes that function alternatively to retain or remove P in the epilimnion throughout the course of a season. The interannual stochasticity of biomass under consistent nutrient inputs could be a result of Lake 227 operating under multiple stable states under high and low N:P loading.

The MyLake model presents several benefits in characterizing the Lake 227 system including detailed physical structuring, nutrient cycling, and sediment diagenesis. Although empirical analysis suggests that top-down controls such as zooplankton grazing do not explain the interannual variation in PP, a model that incorporates many species with different growth kinetics (e.g., Reynolds et al. 2001; Mooij et al. 2007) may be able to better reproduce phytoplankton growth dynamics under varying N:P inputs. However, a community-based modeling approach sacrifices spatiotemporal detail, introducing additional sources of complexity that may overparameterize the model and thereby limit its use. MyLake was successful in predicting lake thermal structure that drives biogeochemical reactions in the system, establishing the capability for scenario testing.

We observed a significant interaction of climate change and fertilization on phytoplankton blooms in Lake 227. Given that nutrient management activities can be partly counteracted and confounded by climate change (Nielsen et al. 2014), evaluating climate- and nutrient-related factors concurrently is a key advance. Experimental P loading is the major driver of blooms in this system, as evidenced by the markedly low PP concentrations under the Climate Change Only scenario. The impacts of catchment and internal loads of P were most strongly observed at the start of the ice-free season (May) in the absence of experimental fertilization. Systems with little anthropogenic P loading may thus observe the effects of climate change most strongly after ice break prior to the establishment of water column stratification. The Climate Change + Fertilization scenario produced larger blooms than the Fertilization Only scenario. Although increases in air temperature resulted in increases in surface water temperature that were an order of magnitude smaller, the timing and persistence of this shift impacted the model’s reaction network such that differences in PP concentrations were substantial. Specifically, the greatest increases in water temperature in the Climate Change + Fertilization scenario occurred in the early part of the ice-free season, allowing blooms to establish earlier and reach concentrations in June, July, and August that were 10.2 ± 6.2 μg/L greater than in the Fertilization Only scenario. We thus demonstrate that warmer surface water temperatures in the spring, in combination with steady P loading, have resulted in higher bloom biomass than would have occurred in the absence of climate change.

This study presents the first efforts to disentangle the effects of nutrient loading regimes and climate change on phytoplankton blooms in Lake 227, a lake unique in its long-term record and history of whole-lake experimentation. The process-oriented model applied in Lake 227 was challenged to reproduce interannual dynamics of phytoplankton blooms associated with different fertilization regimes. Predicting the complex responses of lakes to changing N:P loading is difficult (Filstrup and Downing 2017), and different model structures ought to be tested in order to fully capture the drivers of biomass during high N:P periods. Nonetheless, MyLake performed well in predicting the timing and magnitude of blooms when low N:P ratios favor diazotrophic cyanobacteria, demonstrating the potential to provide insight into HAB mitigation strategies when external nutrient loads are characterized by low N:P. We show that P loading is the dominant driver of phytoplankton blooms in this system, but the concurrent effects of increased temperature and fertilization exacerbate blooms beyond that which would be experienced from fertilization alone.

## Acknowledgements

Thank you to members of the Schiff and Venkiteswaran lab groups for their feedback throughout the project. We are indebted to Richard Elgood for his logistical support and scientific feedback. This project was supported by NSERC Strategic Partnership Grant for Projects (STPGP 494497-2016) and Mitacs Accelerate Cluster Grant (IT10980).

