## Supplemental Information for "Long-term influence of climate and experimental eutrophication regimes on phytoplankton blooms"

Salk et al.

### Methods

*Corrections of meteorological and catchment measurements*

Specifically, wind speeds were scaled by an empirically determined factor of 0.527 to account for the wind sheltering effect of small lake size (Solinske 1982) . Photosynthetically active radiation (PAR) measurements from the meteorological site were scaled by a factor of 2.22 to convert to total radiation (Meek et al. 1984). Gaps in PAR measurements were filled with the average value across the dataset for each Julian day. Inflow temperature was estimated according to the empirical relationship for Minnesota streams presented by (Erickson and Stefan (2000):

$$\text{water temperature } (^{o}\text{C}) = 0.82 \times \text{air temperature} + 4.18 \quad (1)$$

Inflow volume and chemistry from Lake 239 were scaled by the catchment size of Lake 227 following previous examples (e.g., Higgins et al. 2017). Experimental P and N loadings to L227 were added to the natural loads from inflowing streams to construct total nutrient loads. For variables with seasonal trends, gaps between measurements were filled via linear interpolation. For variables with no evident seasonal trends, gaps between measurements were filled with the mean value across the dataset. Dates of ice break and ice freeze were estimated from direct observations from Lake 239 each year, corrected for differences in lake size based on empirical relationships between lake size and ice phenology assessed during 2016 and 2017. The corrected ice break dates for Lake 227 ranged from 3–5 days earlier and corrected ice freeze dates ranged from 9–18 days earlier than dates observed for Lake 239.

*Modified Mann-Kendall for Time Series Analysis*

The Mann-Kendall test is a nonparametric method for identifying the presence of a monotonic trend in time series data. Lake and stream chemistry samples taken biweekly at IISD-ELA are temporally autocorrelated, violating the assumption of independence among samples of the Mann-Kendall test. We thus applied two modifications of the Mann-Kendall test which account for autocorrelation, whereby (1) data are detrended, (2) effective sample size is calculated from the significant serial correlation coefficients, and (3) the variance of the test statistic is modified to remove the effect of autocorrelation. The modification proposed by Hamed and Rao (1998) calculates effective sample size from the correlation coefficients of the ranks of the sample data, but it can overinflate the correction factor and result in a false negative result for the presence of a significant trend (Yue et al. 2002). Yue and Wang (2004) have presented a similar modification that calculates effective sample size from the correlation coefficients of the sample data, avoiding the overestimated rejection rate. The outcomes of the two modified tests were compared with the results of the unmodified Mann-Kendall test (Table S1). The outcome of each test for a single variable was consistent across all three tests, with the exception of TDN:TDP prior to the 1975 change point, where a significant positive trend was found in the Yue and Wang modification and no trend was found in the original Mann-Kendall and Hamed and Rao modification.

*Application of MyLake to Lake 227*

At each daily time step of the MyLake model, the thermal structure of the lake is set up from surface and sediment heat fluxes, vertical diffusion, wind stress, and stream inflow. Chemical and biological reactions take place under a system of differential equations, and the vertical distribution of chemical species are influenced by diffusion and advection terms. The parameters associated with reactions may be adjusted by the user. Light attenuation is governed by the absorption, scattering, and shading of light by water, DOC, and phytoplankton cells. Phytoplankton growth and biomass accrual is determined by P uptake kinetics, light, and temperature and includes loss terms for respiration, sedimentation, and remineralization. Additional details for the MyLake model can be found in the user guide (Saloranta and Andersen 2007)

Application of MyLake to Lake 227 began with a characterization of conditions in 1969. Initial temperature and chemistry profiles of the water column were specified from measurements made immediately prior to the first fertilization event in June 1969. Initial sediment chemistry and redox profiles were determined from sediment core data isolated for the pre-fertilization period (Hesslein 1980; Schindler et al. 1987; Ansems 2012). Subsequent daily time steps of the model required input data for meteorological conditions and the temperature and chemical concentrations of stream inflows to the lake. Parameters for each reaction equation were specified, and the reactions as well as external meteorological and inflow forcings drove daily water column conditions in the lake. Parameter values for reaction equations were based on an application of the model to the humic and eutrophic Lake Vansjø, Norway (Saloranta and Andersen 2007; Couture et al. 2014, 2018). When available, direct measurements from Lake 227 (e.g., C:N:P ratios of phytoplankton) were supplied as parameter values. Parameter values were optimized using the Genetic Algorithm function of the MATLAB Global Optimization Toolbox, which sought to minimize the sum of squared error between observed and modeled PP and TDP by varying parameters within the maximum probable range. Based on sensitivity analysis in previous MyLake applications (Saloranta and Andersen 2007; Couture et al. 2014), PP concentrations are most sensitive to the following parameters: the PAR saturation level for phytoplankton growth, half saturation growth P level, growth rate, loss rate, and sinking rate. For oxygen and DOC, the same optimization algorithm was run for the rate constant for DOC mineralization, the half-saturation coefficient for oxic metabolism, and the inflow scaling factor for DOC concentration.

*Model Fit Metrics*

Aggregated performance metrics for temperature, PP, TDP, and $O_2$ were calculated in terms of normalized bias (*B**) and normalized unbiased root mean squared difference (*RMSD'**) (Los and Blaas 2010):

$$B^{*} = \frac{\bar{M} - \bar{D}}{\sigma_D} \tag{2}$$

$$RMSD'^{*} = \frac{\mathrm{sgn}(\sigma_M - \sigma_D)}{\sigma_D} \left[ \overline{(M'_n - D'_n)^2} \right]^{0.5} \tag{3}$$

Where *M* indicates model output, *D* indicates observations, $\sigma_D$ and $\sigma_M$ are the standard deviations of the observations and model output, respectively, sgn is the sign of the standard deviation difference, and $M'_n$ and $D'_n$ are the residuals of individual modeled and observed values compared to the average modeled and observed values, respectively. The overbar in equations 2 and 3 denotes averaging.

**Figures**

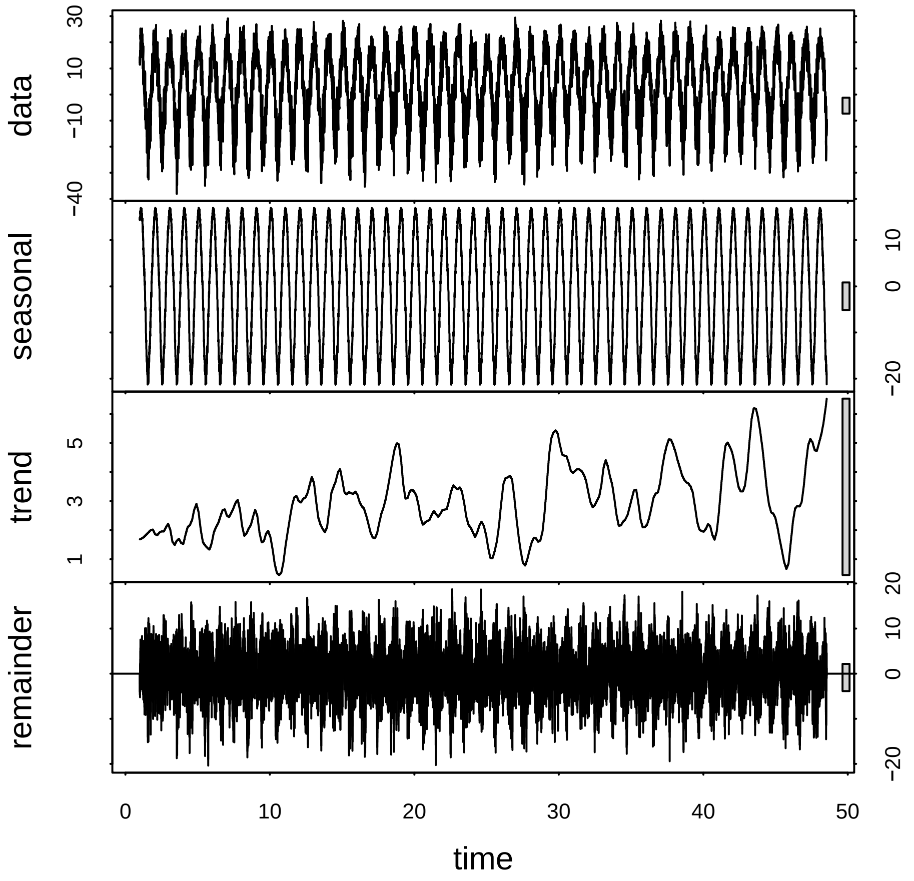


**Figure S1.** Observations in air temperature at Lake 227 (top panel), broken down by seasonal variation (second panel), systematic trend (third panel), and random variation (bottom panel) using loess-based time series decomposition. Y axes are temperature in degrees C (note difference in scales), and x axis is years since 1969.

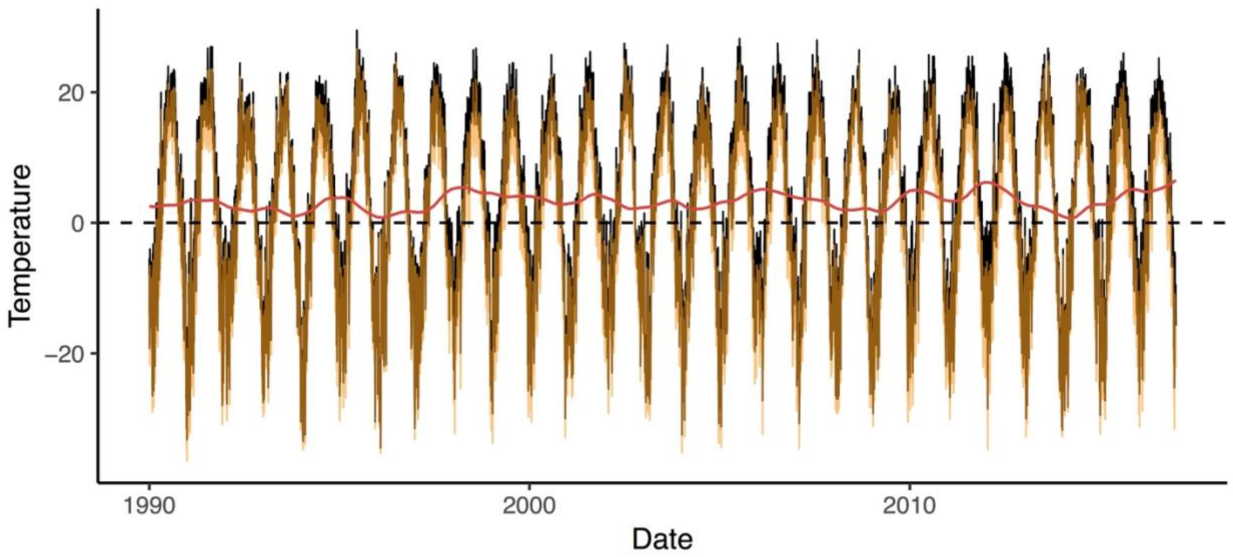


**Figure S2.** Air temperature observed at Lake 227 (black) from 1990-2016, with detrended temperature (yellow) as calculated by the observed temperature minus the systematic trend since 1969 (red). Detrended temperature is 3.2 ± 1.3 ºC warmer than the observed temperature, with a total range of 0.7–6.5 ºC.

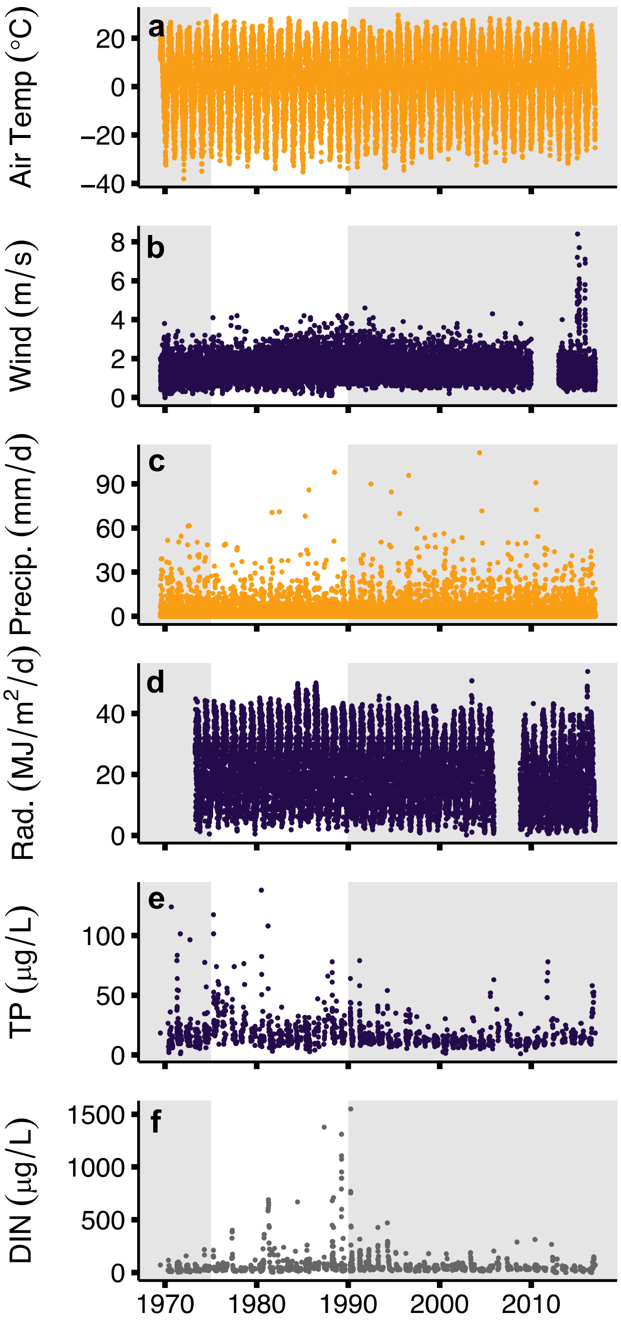


**Figure S3.** Observed historical trends in external drivers in the Lake 227 catchment. (a-d) Conditions at the meteorological site (Precip. = precipitation, Rad. = radiation). (e-f) Stream inflow concentrations of total P and dissolved inorganic N. Shading represents fertilization period. Color of point indicates direction of trend as determined by seasonal Mann-Kendall test (a-d) and Mann-Kendall test (e-f) (yellow = significant positive trend, purple = significant negative trend, gray = non-significant trend).

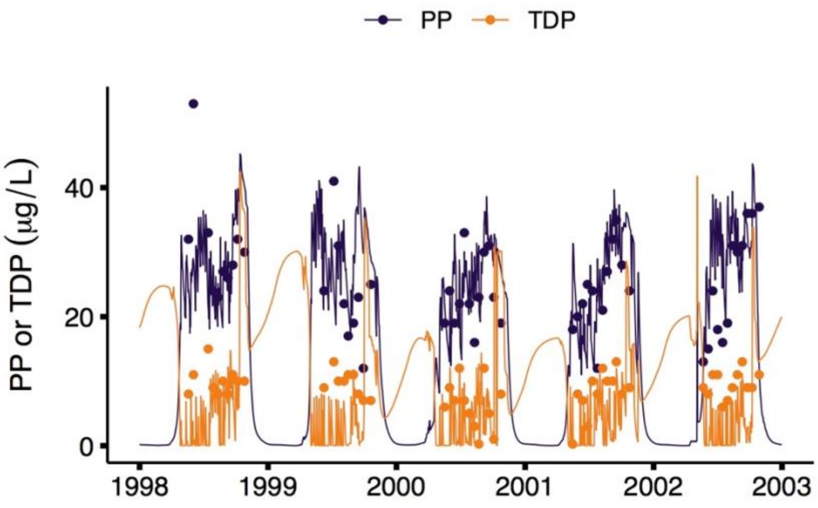


**Figure S4.** Observed (points) and modeled (lines) epilimnetic PP and TDP concentrations for the five best model fit years, as evaluated by least squares regression analysis.

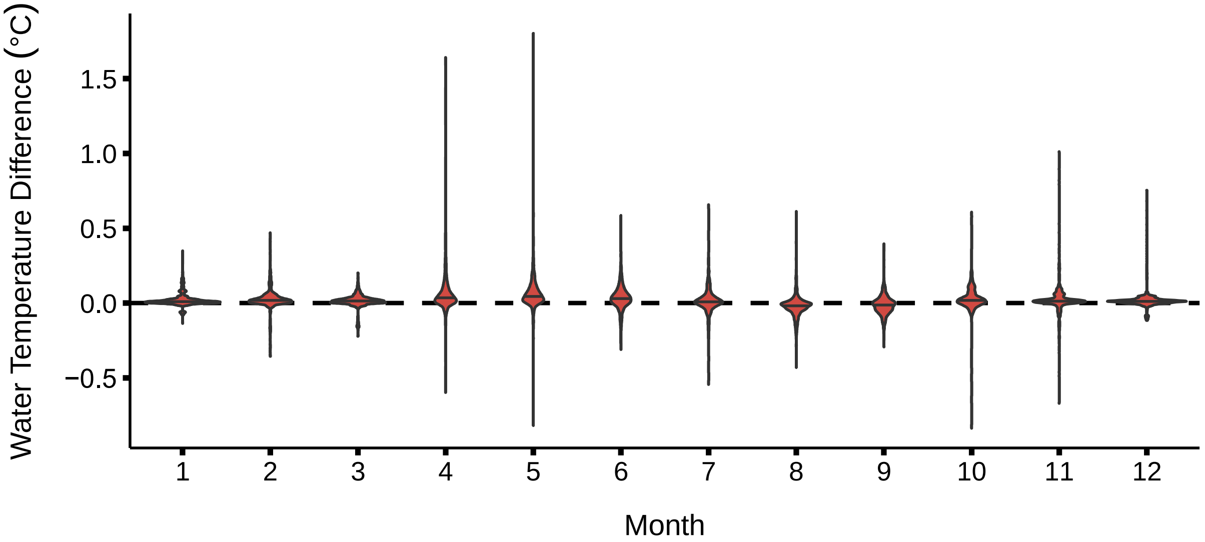


**Figure S5.** The difference in daily temperature measurements at 1 m depth between the Climate Change + Fertilization and Fertilization Only scenarios, pooled for each month across modeled years (1990-2016). Violins represent the total range in values along the y axis, with a density function represented on the x axis. Positive values indicate higher temperature in the Climate Change + Fertilization scenario than in the Fertilization Only scenario.

### Tables

**Table S1.** Results of Mann-Kendall tests for monotonic trends in lake and stream data over time. Results are reported for periods before and after a change point was detected via Pettitt's test. For variables where a change point was identified, results before and after the change point are reported as the upper and lower values, respectively. z = z-score, p = p-value, N/A = not assessed.

| Variable | Change Point | Sen's Slope | Modified M-K (Yue & Wang 2004) | | Modified M-K (Hamed & Rao 1998) | | Mann-Kendall | |
|---|---|---|---|---|---|---|---|---|
| | | | z | p | z | p | z | p |
| TN:TP (molar) | 1996 | 0.027 | **6.05** | **< 0.0001** | **4.95** | **< 0.0001** | **4.02** | **< 0.0001** |
| | | 0.007 | 1.69 | 0.09 | 0.68 | 0.49 | 0.77 | 0.44 |
| TDN:TDP (molar) | 1975 | 0.182 | **2.47** | **0.01** | 0.95 | 0.34 | 1.44 | 0.15 |
| | | -0.032 | **-5.08** | **< 0.0001** | **-1.97** | **0.049** | **-3.53** | **< 0.001** |
| PP (μg/L) | 1978 | 0.021 | 1.22 | 0.22 | 0.62 | 0.53 | 1.32 | 0.19 |
| | | -0.019 | **-3.89** | **< 0.001** | **-3.33** | **< 0.001** | **-6.78** | **< 0.0001** |
| chl (μg/L) | 1987 | -0.023 | **-3.45** | **< 0.001** | **-2.08** | **0.037** | **-5.02** | **< 0.0001** |
| | | -0.020 | **-3.59** | **< 0.001** | **-2.01** | **0.044** | **-3.71** | **< 0.001** |
| Inflow TP | N/A | -0.005 | **-5.07** | **< 0.0001** | **-3.81** | **< 0.001** | **-9.35** | **< 0.0001** |
| Inflow DIN | N/A | 0.001 | 0.15 | 0.88 | 0.10 | 0.92 | 0.34 | 0.74 |

**Table S2.** Results of Seasonal Mann-Kendall tests for monotonic trends in meteorological data over time. z = z-score, p = p-value.

| | z | p |
|---|---|---|
| Air temp (°C) | 7.23 | <0.0001 |
| Wind speed (m/s) | -2.72 | <0.01 |
| Precip. (mm/d) | 8.86 | <0.0001 |
| Radiation (MJ/m$^2$/d) | -23.29 | <0.0001 |

**Table S3.** Root-mean squared error (RMSE) values, representing the average deviation of modeled values from observed values in the units of interest, for three fertilization periods in Lake 227.

| | High N:P | Low N:P | P-only | Units |
|---|---|---|---|---|
| Ice Break | 1.35 | 3.21 | 5.92 | d |
| Ice Freeze | 1.05 | 4.93 | 5.65 | d |
| Temp 1 m | 1.75 | 1.37 | 2.03 | °C |
| Temp 4 m | 1.28 | 1.78 | 1.62 | °C |
| Temp 9 m | 0.31 | 0.33 | 0.58 | °C |
| PP | 20.63 | 21.76 | 10.49 | μg $L^{-1}$ |
| TDP | 4.91 | 3.62 | 3.69 | μg $L^{-1}$ |

**Table S4.** PP- and TDP-sensitive parameters derived from model optimization routines. The first five years of each period were used as optimization periods. PAR_sat = PAR saturation level for phytoplankton growth (mol(quanta) $m^{-2}$ $s^{-1}$), w_chl = settling velocity for phytoplankton (m $d^{-1}$), m_twty = loss rate of phytoplankton ($d^{-1}$) at 20 °C, g_twty = specific growth rate of phytoplankton ($d^{-1}$) at 20 °C, P_half = half saturation growth P level (mg $m^{-3}$).

| Parameter | High N:P | Low N:P | P-only |
|---|---|---|---|
| PAR_sat | 2.38e-4 | 2.84e-4 | 2.46 e-4 |
| w_chl | 0.0978 | 0.0934 | 0.0785 |
| m_twty | 0.0104 | 0.01 | 0.01 |
| g_twty | 0.614 | 0.612 | 0.656 |
| P_half | 3.42 | 0.233 | 0.483 |
